# Factors Associated with Emerging and Re-emerging of SARS-CoV-2 Variants

**DOI:** 10.1101/2021.03.24.436850

**Authors:** Austin N. Spratt, Saathvik R. Kannan, Lucas T. Woods, Gary A. Weisman, Thomas P. Quinn, Christian L. Lorson, Anders Sönnerborg, Siddappa N. Byrareddy, Kamal Singh

**Affiliations:** Christopher S. Bond Life Sciences Center, University of Missouri, Columbia, MO 65211, USA; Department of Biochemistry, University of Missouri, Columbia, MO 65211, USA; Department of Veterinary Pathobiology, University of Missouri, Columbia, MO 65211, USA; Division of Infectious Diseases, Department of Medicine, Karolinska Institute, Huddinge 14186 Stockholm, Sweden; Division of Clinical Microbiology, Department of Laboratory Medicine, Karolinska Institute, Huddinge 14186, Stockholm, Sweden; Department of Pharmacology and Experimental Neuroscience, University of Nebraska Medical Centre, Omaha, NE 68198, USA; Sanctum Therapeutics Corporation, Sunnyvale, CA 94087, USA

**Keywords:** SARS-CoV-2, SARS-CoV, COVID-19, B.1.1.7, B.1.351, P.1, CAL.20C

## Abstract

Global spread of Severe Acute Respiratory Syndrome Coronavirus-2 (SARS-CoV-2) has triggered unprecedented scientific efforts, as well as containment and treatment measures. Despite these efforts, SARS-CoV-2 infections remain unmanageable in some parts of the world. Due to inherent mutability of RNA viruses, it is not surprising that the SARS-CoV-2 genome has been continuously evolving since its emergence. Recently, four functionally distinct variants, B.1.1.7, B.1.351, P.1 and CAL.20C, have been identified, and they appear to more infectious and transmissible than the original (Wuhan-Hu-1) virus. Here we provide evidence based upon a combination of bioinformatics and structural approaches that can explain the higher infectivity of the new variants. Our results show that the greater infectivity of SARS-CoV-2 than SARS-CoV can be attributed to a combination of several factors, including alternate receptors. Additionally, we show that new SARS-CoV-2 variants emerged in the background of D614G in Spike protein and P323L in RNA polymerase. The correlation analyses showed that all mutations in specific variants did not evolve simultaneously. Instead, some mutations evolved most likely to compensate for the viral fitness.

## 1. Introduction

Severe Acute Respiratory Syndrome Coronavirus-2 (SARS-CoV-2), the virus responsible for Coronavirus Disease 2019 (COVID-19), is the seventh human coronavirus (hCoV) discovered to date. The other six are hCoV-229E, hCoV-NL63, hCoV-HKU1, hCoV-OC43, SARS-CoV and Middle East Respiratory Syndrome (MERS) CoV. Four CoVs (hCoV-229E, hCoV-NL63, hCoV-HKU1 and hCoV-OC43) typically cause mild self-limiting cold-like diseases. However, three CoVs (SARS-CoV, MERS-CoV and SARS-CoV-2) cause fatal respiratory illness. CoVs are single-stranded, non-segmented positive (+)-sense RNA viruses of ∼30 kb genome length. They encode up to 16 nonstructural proteins (nsps) and 10-12 structural/accessory proteins [1,2]. The nsps form the replication/transcription complex (RTC) that synthesizes the nascent (-) strand RNA to be used as a template for (+) sense genome and a set of subgenomic RNAs (sgRNA) with common 5’ -leader sequence and a 3’ -poly-A tail. Of the 10-12 structural proteins, there are four major proteins: spike (S), nucleocapsid (N), membrane (M) and envelope (E) that participate in the infection and assembly of infectious progeny virus [3,4].

In the first step of viral infection, the spike (S) protein of hCoVs directly interacts with distinct host cell receptors, where angiotensin-converting enzyme 2 (ACE2) serves as the cell-surface receptor for SARS-CoV and SARS-CoV-2. The synergistic action of host cell serine protease TMPRSS2 to the ACE2/spike interaction cleaves S protein into S1 and S2 subunits to facilitate endocytosis of viral particle [5]. Due to its primary role in hCoV infections, S protein is a crucial target for developing vaccines, therapeutic antibodies, and diagnostics [6]. Structural data show that the receptor-binding domains (RBDs) within the S1 subunits of both SARS-CoV and SARS-CoV-2 have largely conserved interactions with the ACE2, despite being the domain of lowest homology between the two viruses [7-10].

Owing to the critical role of the RBD in SARS-CoV-2 infection, almost all vaccines in development target the RBD. Since currently approved vaccines are based on the RBD sequence of the first isolated virus (Wuhan-Hu-1; GenBank accession no. NC_045512), newly emerged SARS-CoV-2 variants containing mutations within the RBD may present a challenge to vaccination efforts. Here, we present the results of our analyses and show that the combination of three mutations D614G, P323L and C241U are limited to the U. S., and the major SARS-CoV-2 variants (B.1.1.7, the UK variant; B.1.351, the South Africa variant; P.1, the Brazil variant and CAL.20C, the California variant) emerged independently. We then mapped the mutations onto the tertiary and quaternary structural data to evaluate the factors that may be responsible for the apparent enhanced infectivity of the newly emerged SARS-CoV-2 variants. Our results show that (i) N501Y, the most common mutation among all variants, forms a π-π interaction with Y44 of ACE2 that may enhance the binding affinity of S protein with ACE2 and (ii) the mutations in the virus change the conformational entropy of the S protein and thereby its stability. We also show that mutations near these sites may enhance the propensity of virus interaction with certain integrins, presumably alternate/additional receptors of the SARS-CoV-2.

## 2. Results

### 2.1. A comparison of SARS-CoV and SARS-CoV-2 S proteins interaction with ACE2

Three CoVs (hCoV-NL63, SARS-CoV and SARS-CoV-2) use ACE2 as a host cell receptor. The S protein RBDs (S-RBDs) and ACE2 complex have been solved [9-14], which show that the S-RBDs of SARS-CoV and SARS-CoV-2 share structural homology, whereas the S-RBD of hCoV-NL63 is structurally distinct. Nonetheless, the S-RBDs of the three CoVs bind to overlapping sites on ACE2 [11]. To date, three crystal structures [9,12,13] and one cryoEM structure of the S-RBD/ACE2 complex [14] have been reported, as well as a series of cryoEM structures representing different conformations and stoichiometries of the S protein/ACE2 complexes [15]. Two crystal structures of the S protein/ACE2 complexes contain S-RBDs that are identical to the S-RBD of the Wuhan-Hu-1 isolate [9,12], whereas the third crystal structure contains RBD modifications at ∼20 sites [13] (**Fig. 1a**). Although some of these modifications (V417K and N439R) are at the S-RBD/ACE2 interface, they do not appear to affect overall binding interactions as noted in other S-RBD/ACE2 complexes. The cryoEM structure complexes containing one S protein and one, two or three ACE2 proteins provide a great deal of atomic details of spike/ACE2 interactions and suggest that more than one ACE2 can bind to the trimer of the S protein. This is the first report suggesting a higher-order interaction between the two proteins [15]. A total of 14 RBD residues of SARS-CoV and SARS-CoV-2 interact with ACE2 (**Fig. 1a**, red/pink-shaded columns) [16]. Overall, there exists ∼76% homology between S proteins of SARS-CoV and SARS-CoV-2. The most variable region of S protein is the RBD, and the most variable region within the RBD is the receptor-binding motif (RBM) (**Fig. 1a**; green-shaded regions). Despite these variations, 8 out of 14 residues are topologically conserved between SARS-CoV and SARS-CoV-2 (red-shaded residues in **Fig. 1a**) and the remaining 6 residues form chemically similar interactions. Notably, the structural comparison does not reveal dramatic differences in S-RBD residue interactions with ACE2 that can explain the greater binding affinity for ACE2 of the S-RBD of SARS-CoV-2 compared to SARS-CoV.

**Figure 1.**
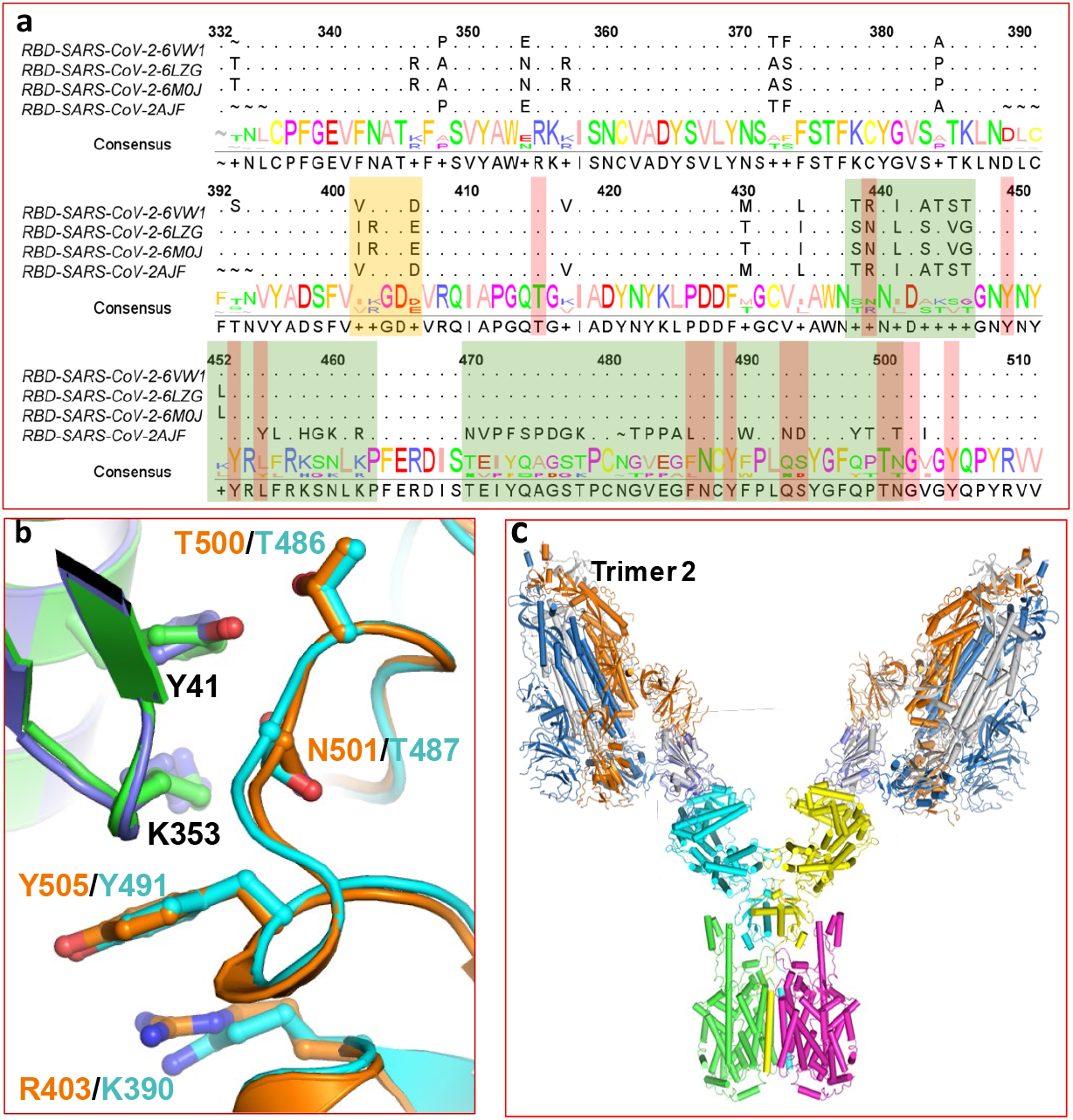
Details of genetics and structural components likely to contribute to SARS-CoV-2 binding, entry and infectivity. **a**. Structure-based sequence alignment of S-RBDs from SARS-CoV (PDB entry 2AJF) and SARS-CoV-2 (Protein Data Bank entries 6WV1, 6LZG and 6M0J). Green-shaded regions represent the receptor-binding motif (RBM), the orange-shaded region represents the RGD motif, and the red/pink-shaded residues in the RBD directly interact with ACE2. **b**. Location of RGD and KGD motifs in S-RBDs of SARS-CoV-2 (orange ribbon) and SARS-CoV (cyan ribbon) and the KGD motif in ACE2 (violet and green ribbons). Residues are shown in ball-and-stick representation with carbons in orange for SARS-CoV-2; cyan for SARS-CoV, and green/violet for ACE2. Oxygen and nitrogen atoms are shown in red and blue colors, respectively. Y41 and K353 belong to ACE2. **c**. A proposed model for binding two S trimers with ACE2 based on the SARS-CoV-2 S-RBD/B^0^AT1/ACE2 complex structure. The S-RBDs of the 2 trimers were superposed on the S-RBD in the cryoEM structure of the S-RBD/B^0^AT1/ACE2 complex.

Several structural studies also experimentally determined the binding affinity between S-RBD and ACE2. Using Biolayer Interferometry (BLI), Walls *et al*. showed that S-RBD of SARS-CoV-2 binds to ACE2 with nearly a 4-fold greater affinity than S-RBD of SARS-CoV [16]. Surface plasmon resonance (SPR) measurements have shown an ∼4-fold [13] to ∼20-fold [8] greater binding affinity of ACE2 with SARS-CoV-2 S-RBD compared to SARS-CoV S-RBD. In contrast, little difference between the binding affinities of SARS-CoV and SARS-CoV-2 S-RBD for ACE2 was reported elsewhere [9]. Therefore, it remains unclear if differences in the binding affinities of S-RBDs for ACE2 are responsible for the greater infectivity of SARS-CoV-2 compared to SARS-CoV. Thus, we explored alternative mechanisms to explain the higher infectivity of SARS-CoV-2, focusing on the mechanisms of SARS-CoV-2 entry after S protein binding to ACE2.

### 2.2. Alternate SARS-CoV-2 entry mechanisms

A notable difference between SARS-CoV-2 and other human CoVs is the presence of a conserved Arg-Gly-Asp (RGD) motif (403-405) in the S-RBD [17]. The same position in SARS-CoV S-RBD is occupied by a Lys-Gly-Asp (KGD) motif that is functionally related to an RGD motif (**Fig. 1a**, orange-shaded region) [18]. In the family of ∼20 heterodimeric integrin molecules, 9 RGD-binding integrins can directly interact with RGD motifs of extracellular matrix proteins to promote normal tissue morphology and regulate physiological functions cell adhesion and migration [19,20]. RGD-binding integrins, when activated by RGD-containing ligands, are also able to modulate outside-in signaling under a wide variety of conditions [19,21-23]. Integrins are known to be utilized by certain viruses to bind the host cells [24]. Thus, several human viruses, including metapneumonovirus [25], human cytomegalovirus [26] and adenovirus types 2 and 5 [27] use RGD motifs for attachment to RGD-binding integrins. An examination of the structures of SARS-CoV-2 S-RBD/ACE2 complexes indicates that the RGD motif in the S-RBD does not directly interact with ACE2, despite its location near the middle of the S-RBD/ACE2 interface, suggesting an alternate role for the RGD motif of the SARS-CoV-2 S-RBD. It is postulated that RGD-binding integrins recognize the RGD motif of SARS-CoV-2 S-RBD and that these integrins [28] could serve as an alternate SARS-CoV-2 receptor or as a co-receptor together with ACE2, thereby contributing to the higher infectivity of SARS-CoV-2 compared to SARS-CoV.

ACE2 also contains both RGD (203-205) and KGD (353-355) motifs and has been shown to bind integrins and modulate outside-in signaling [29]. RGD 203-205 is distal to the bound S-RBD of SARS-CoV-2 and is near the active site cleft formed by two domains of ACE2 and, therefore it should not be accessible for protein-protein interaction. Indeed, ACE2 interaction with α5β1 integrins was found to be independent of this RGD motif. However, the KGD 353-355 motif of ACE2 is localized at the S-RBD interface (**Fig. 1b)**, recognized by RGD-binding integrins [18], and mutation in this motif strongly inhibits binding of SARS-CoV S protein [30]. Residues R403 of the RGD motif in the S protein of SARS-CoV-2 (KGD in SARS-CoV) and K353 of the KGD motif of ACE2 are oriented in such a way that they sandwich Y505 of SARS-CoV-2 S-RBD (Y491 for SARS-CoV) (**Fig. 1b**), which in turn forms conserved interactions with E37 and R393 of ACE2 (not shown). Additionally, the RGD motif of S-RBD in SARS-CoV-2 and the KGD motif of SARS-CoV as well as the KGD motif in ACE2 appear to be strategically positioned near Y41 of ACE2, which interacts with N501 and T500 of SARS-CoV-2 or T487 and T486 of SARS-CoV (**Fig. 1b**). Taken together, this analysis implies that SARS-CoV-2/SARS-CoV may have evolved an S protein RGD/KGD motif that enhances association with host cells through RGD-binding integrins to optimize cell entry of SARS-CoV/ACE2 complexes.

### 2.3. Binding of two S protein trimers to the ACE2 dimer

A cryoEM structure of SARS-CoV-2 S-RBD, ACE2 and sodium-dependent neutral amino acid transporter B^0^AT1 (SLC6A19) showed that ACE2 in the dimeric form interacts with SLC6A19, and two SARS-CoV-2 S-RBDs bind to the ACE2 dimer [14]. For functional expression on the cell surface, SLC6A19 requires an obligatory subunit, collectrin (TMEM27) in the kidney and ACE2 in the intestine [31-34]. The binding interface for the two S-RBDs on ACE2 is the same as reported for the crystal structures, as mentioned earlier. However, this report suggests that two S protein trimers can bind to the ACE2 dimer (**Fig. 1c**) when bound to B^0^AT1, thereby increasing the infectivity of SARS-CoV-2. However, the structure of the SARS-CoV S-RBD/B^0^AT1/ACE2 complex is not known and it remains to be seen whether these S protein trimers can enhance the infectivity of SARS-CoV-2.

### 2.4. Geo-prevalence of re-emerging SARS-CoV-2

Several new SARS-CoV-2 variants have emerged over the past few months. The four major variants are characterized by a set of mutations in the S protein, and a few mutations in different ORFs (**Supplementary Text**). A list of the mutations in the four major variants is given in **Table 1**. To understand if SARS-CoV-2 S proteins containing these mutations (**Table 1**) vary regionally or co-exist in different variants, we analyzed sequences (n = 7, 232) from different regions: Brazil (n = 119), California (n = 1,683), South Africa (n = 129) and the United Kingdom (n = 5,301). The results presented in **Fig. 2** show that the S protein mutations among new variants are limited almost exclusively to specific variants and geographic locations. There are only two exceptions. D614G is present universally at high (∼100%) prevalence and N501Y is present in P.1, CAL.20C, B.1.351 and B.1.1.7 at 56, 6, 99 and 83% frequencies, respectively (**Fig. 2**). The low prevalence of common mutations among different variants also suggests that these variants have evolved independently. All regions also had other mutations in addition to their respective variant’ s signature mutations. For example, the CAL.20C variant is characterized by four mutations (S13I, W152C, L452R, D614G). However, mutations P26S, N501Y, A570D, P681H, S982A and D1118H were significantly prevalent (up to 9%) in California. Additionally, the V1176F mutation, which has not been included as a signature mutation of the P.1 variant, was present at a very high frequency (∼81%). It is worth mentioning here that the N501Y mutation that is a signature mutation of the UK (B.1.17), Brazil (P.1) and SA (B.1.351) variants is significantly less prevalent (∼6%) in the CAL.20C variant. The data presented in **Fig. 2** also indicate the prevalence of different variants in the USA (except CAL.20C, discussed in Section 2.7). Thus, considering that P681H is a signature mutation of B.1.1.7, nearly 9% of infections in the USA (*i*.*e*. CAL.20C) may be related to B.1.1.7 (**Fig. 2**).

**Table 1:**
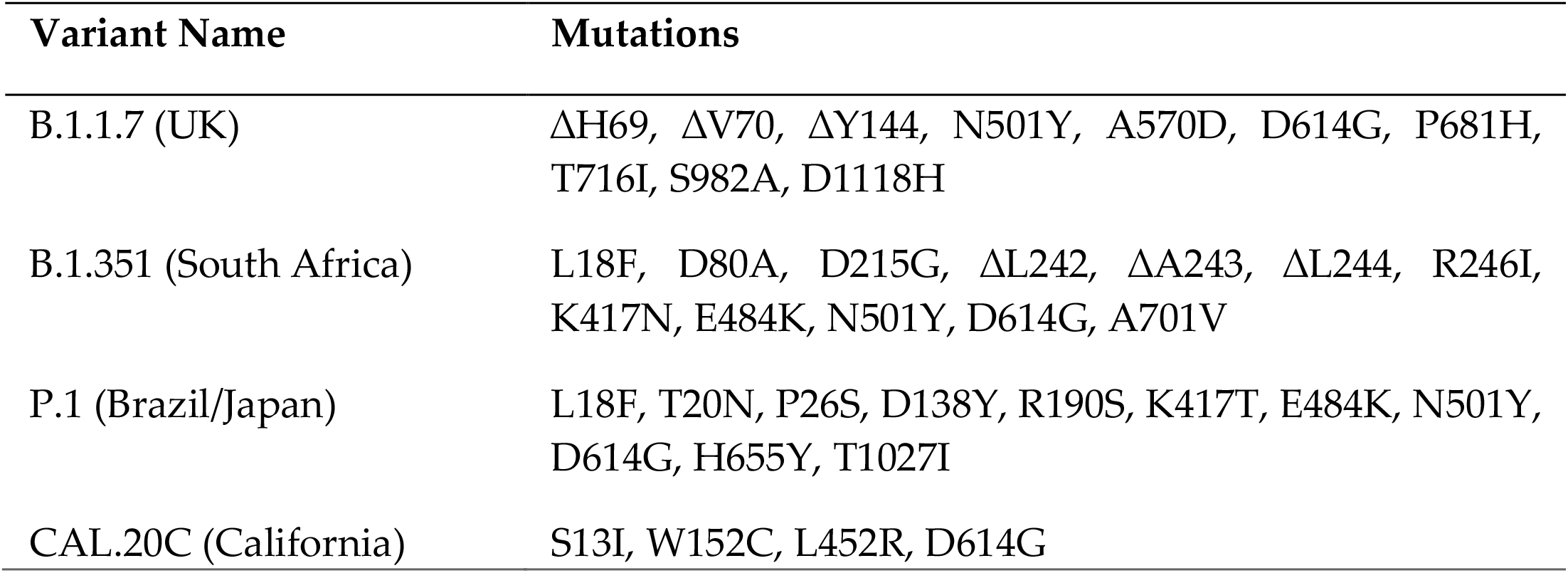
Major new SARS-CoV-2 variants and mutations in the variants.

| Variant Name | Mutations |
| --- | --- |
| B.1.1.7 (UK) | ΔH69, ΔV70, ΔY144, N501Y, A570D, D614G, P681H, T716I, S982A, D1118H |
| B.1.351 (South Africa) | L18F, D80A, D215G, ΔL242, ΔA243, ΔL244, R246I, K417N, E484K, N501Y, D614G, A701V |
| P.1 (Brazil/Japan) | L18F, T20N, P26S, D138Y, R190S, K417T, E484K, N501Y, D614G, H655Y, T1027I |
| CAL.20C (California) | S13I, W152C, L452R, D614G |

**Figure 2.**
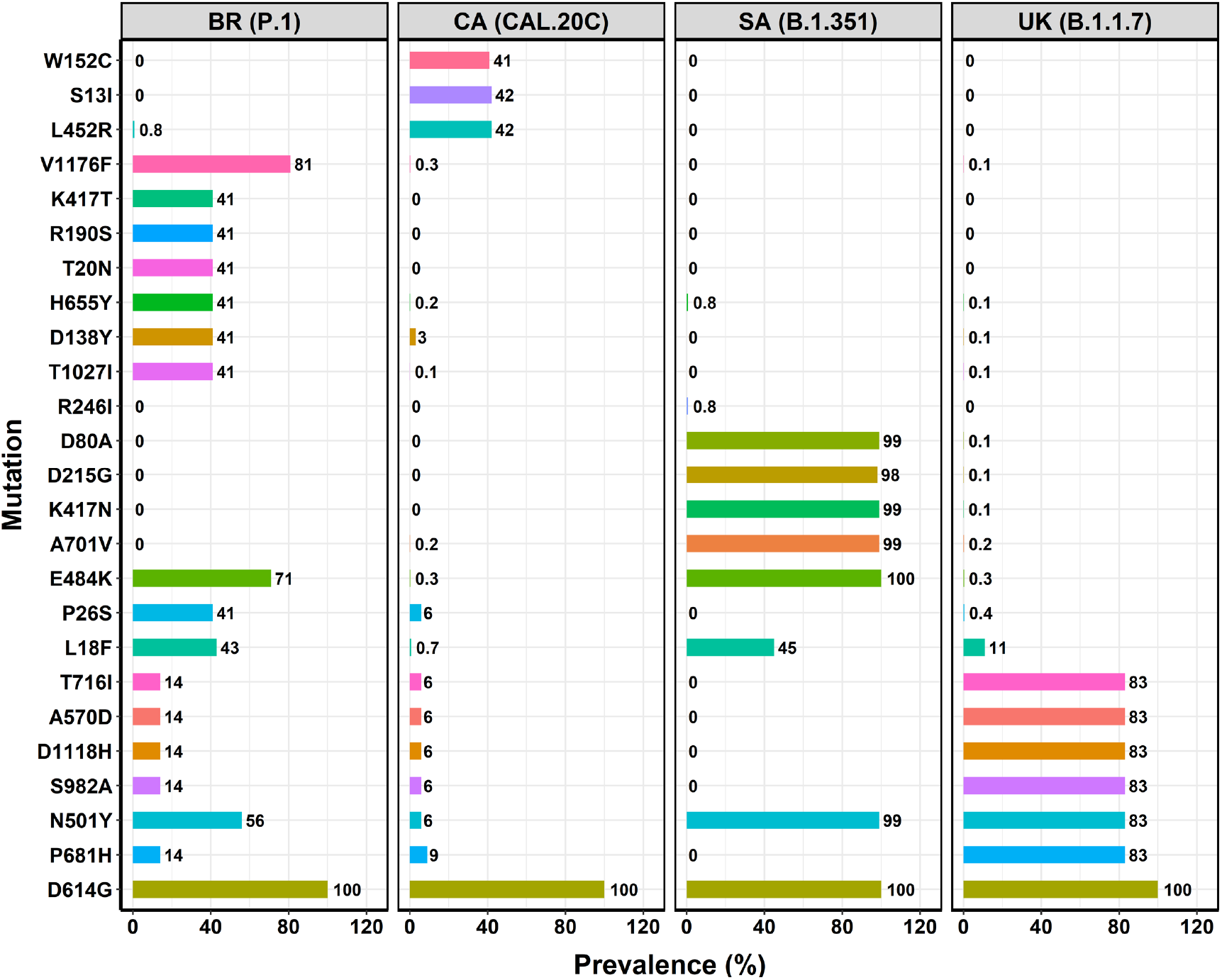
Geo-prevalence of mutations within S protein of four variants. Additional mutations were added to this figure after we noted that these mutations were not reported previously for these variants. For example, the CAL.20C variant has been characterized by S13I, W152C, L452R and D614G mutations. However, we noted additional mutations in this variant (*e. g*. P681H ∼ 9%). Also, previous characterization of the P.1 variant did not include V1176F. Our results showed this mutation was at high prevalence (∼81%). Therefore, this mutation is included in the figure.

### 2.5. Relative abundance of mutations in different variants

To find out the relative abundance of specific mutations within a given variant and with respect to previously reported three co-existing mutations (D614G, P323L and C241U) in the United States [35], we analyzed a significantly larger number of sequences (n = 225,368) that included UK (207,088), South Africa (3,273), Brazil (2,541) and California (12,466). Of these, 47,078 (∼23%) sequences from the UK are B.1.1.7, 590 (∼18%) in South Africa are B.1.351, 61 (∼2%) in Brazil are P.1, and 1,212 (∼10%) in the United States are CAL.20C (as of Feb. 6, 2021). All sequences included in this study have the D614G mutation in the S protein. The results of this analysis are shown in **Supplementary Figure 1**. This figure confirms our two previous observations: (i) three mutations (D614G, P323L and C241U) co-exist in the United States [35], and (ii) variant-specific mutations evolved independently (as concluded from a smaller dataset **Fig. 2**). The analysis of a large number of sequences revealed that all variants had additional mutations than the reported signature mutations from these variants. For example, the prevalence of P681H was 14% in P.1 and 9% in CAL.20C (**Fig. 2**). In addition, the V1176F mutation at a prevalence of 81% co-exists with signature mutation E484K in P.1 (**Fig. 2**).

### 2.6. Correlation among variant-specific mutations

To determine the correlation among variant-specific mutations, we conducted a correlation analysis that included mutations P323L and C241U in addition to the mutations in the S protein. The results presented in **Fig. 3a-d** show that D614G and P323L are highly correlated independent of geographical location in all variants. However, a strong correlation among three mutations (D614G, P323L and C241U) was seen only in the U. S. infections, suggesting that the SARS-CoV-2 virus in the U. S. has been different from the ones in the UK, Brazil and South Africa.

**Figure 3.**
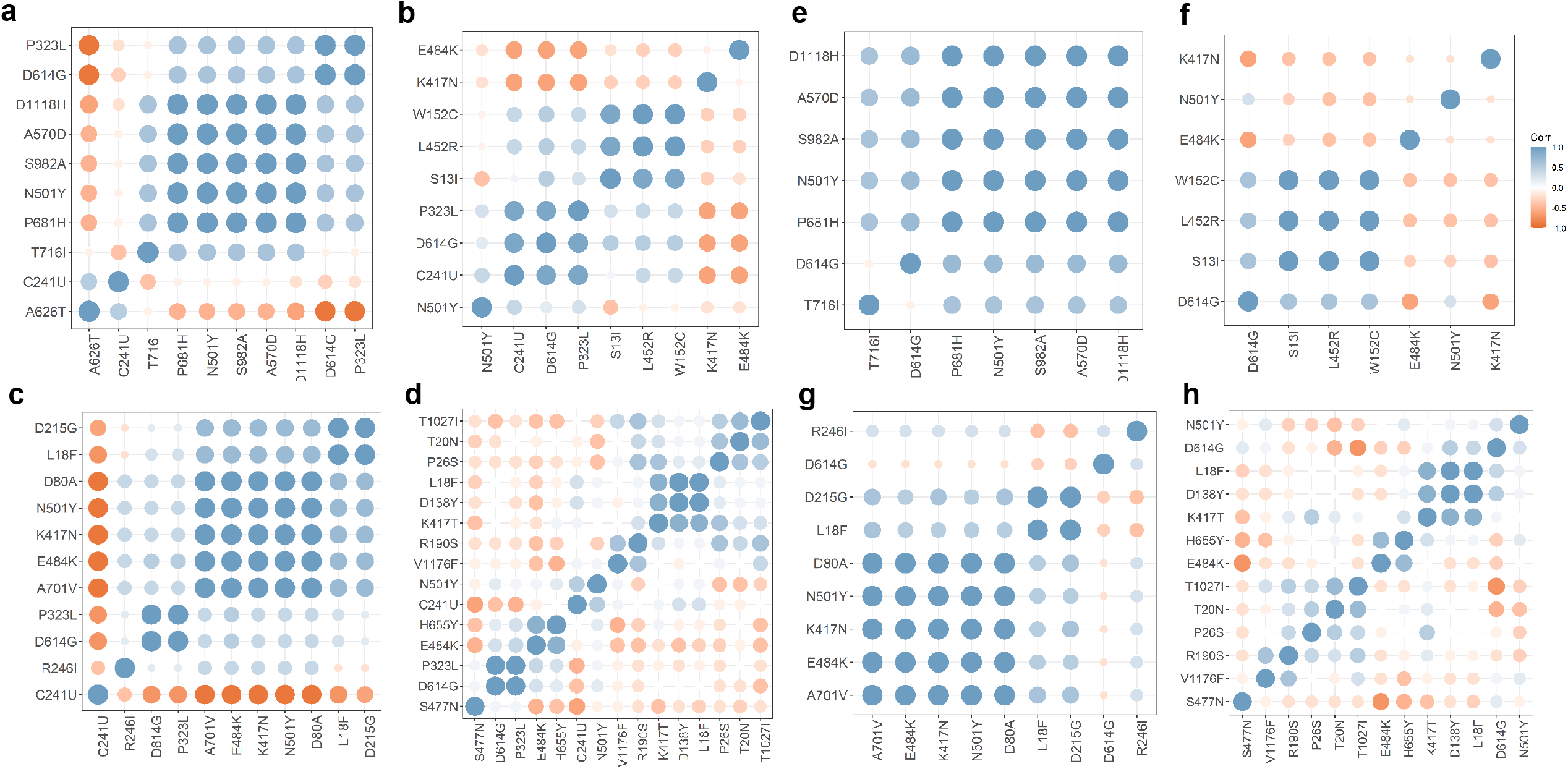
Correlation analyses of mutations in four variants. Panels **a, b, c and d** show the analyses of mutations in S protein together with D614G, P323L and C241U(T) from B.1.1.7, CAL.20C, B.1.351 and P.1 variants, respectively. Panels **e, f, g and h** show the correlation analyses of mutations in S protein from B.1.1.7, CAL.20C, B.1.351, and P.1 variants, respectively. Some additional mutations, V1176F in P.1 and N501Y, E484K and K417N in CAL.20C, were included in our analyses as these mutations were significantly prevalent in respective variants.

To determine the correlation among variant-specific mutations in S protein, we excluded the P323L and C241U mutations. Thus, B.1.1.7 mutations are strongly correlated with D614G except for T716I (**Fig. 3e**), which is negatively correlated (−0.3), suggesting that T716I most likely evolved separately in the background of (D614G, P681H, N501Y, S982A, A570D and D1118H). We also included mutation A626T in our analyses as it is significantly prevalent in the UK viruses (75%) (**Supplementary Fig. S1**) that also contained P323L (**Fig. 3a**).

The CAL.20C (**Fig. 3b**) variant has two distinct groups of correlated mutations. The first group includes D614G, P323L and C241U (**Fig. 3b**), confirming our previous results [35]. The second group of correlated mutations includes S13I, W152C and L452R (**Fig. 3b**). The CAL.20C variant has four signature mutations (S13I, W152C, L452R and D614G) in the S protein (**Table 1 and Fig. 2**). To determine the correlation among the S protein mutations of CAL.20C, we excluded P323L and C241U mutations from the correlation analyses. The results in **Fig. 3f** clearly show that all four S protein mutations are strongly correlated. We also included additional S protein mutations in our analyses based on data presented in **Fig. 2**. The results show a positive correlation of N501Y with D614G, suggesting that these mutations coexist in some viruses in the United States. It is possible that another variant in the U. S. may emerge that includes D614G and N501Y mutations.

As expected, all the mutations in B.1.351 (SA variant) are correlated except D614G, which clustered with P323L (**Fig. 3c**). To gain insight into the correlation among S protein mutations, we re-analyzed the SA sequences after excluding P323L and C241U mutations. The correlation analysis presented in **Fig. 3g** reveals that mutations in the S protein of the B.1.351 variant are positively correlated except for D614G and R246I, which are clustered together. These results indicate a complex evolution of the mutations in B.1.351 S protein. Considering that a majority of viruses contain D614G, it appears that R246I emerged separately than other B.1.351 mutations.

The correlation analysis of the mutations in the P.1 (Brazil) variant is extremely complex (**Fig. 3d**). There are at least 6 groups of correlated mutations. While D614G and P323L are positively correlated, C241U does not cluster with D614G and P323L. Instead, C241U shows a positive correlation with N501Y. To better understand the correlation of mutations in the S protein of P.1, we excluded P323L and C241U and re-analyzed the correlation of mutations (**Fig. 3h**). There are two major groups of positively correlated mutations. The first group includes D614G, N501Y, L18F, D138Y and K417T, whereas the second group of correlated mutations includes T20N, P26S, R190S and T1027I. Additionally, the V1176F mutation is positively correlated with R190S. Considering that V1176F is not a signature mutation in the P.1 variant, a different and probably more complex variant may emerge from P.1.

### 2.7. Prevalence of CAL.20C in different states of the United States

We used signature mutations of CAL.20C to map the prevalence of this variant in the United States. As of Feb. 24, 2021, 36 states in the U. S. had CAL.20C infections. A map of the CAL.20C distribution in the USA is shown in **Fig. 4**. Since no sequences from South Dakota have been deposited, it was not possible to determine the prevalence of CAL.20C there. Specific details of mutation prevalence of four signature S protein mutations in CAL.20C are shown in **Supplementary Figure 2**.

**Figure 4.**
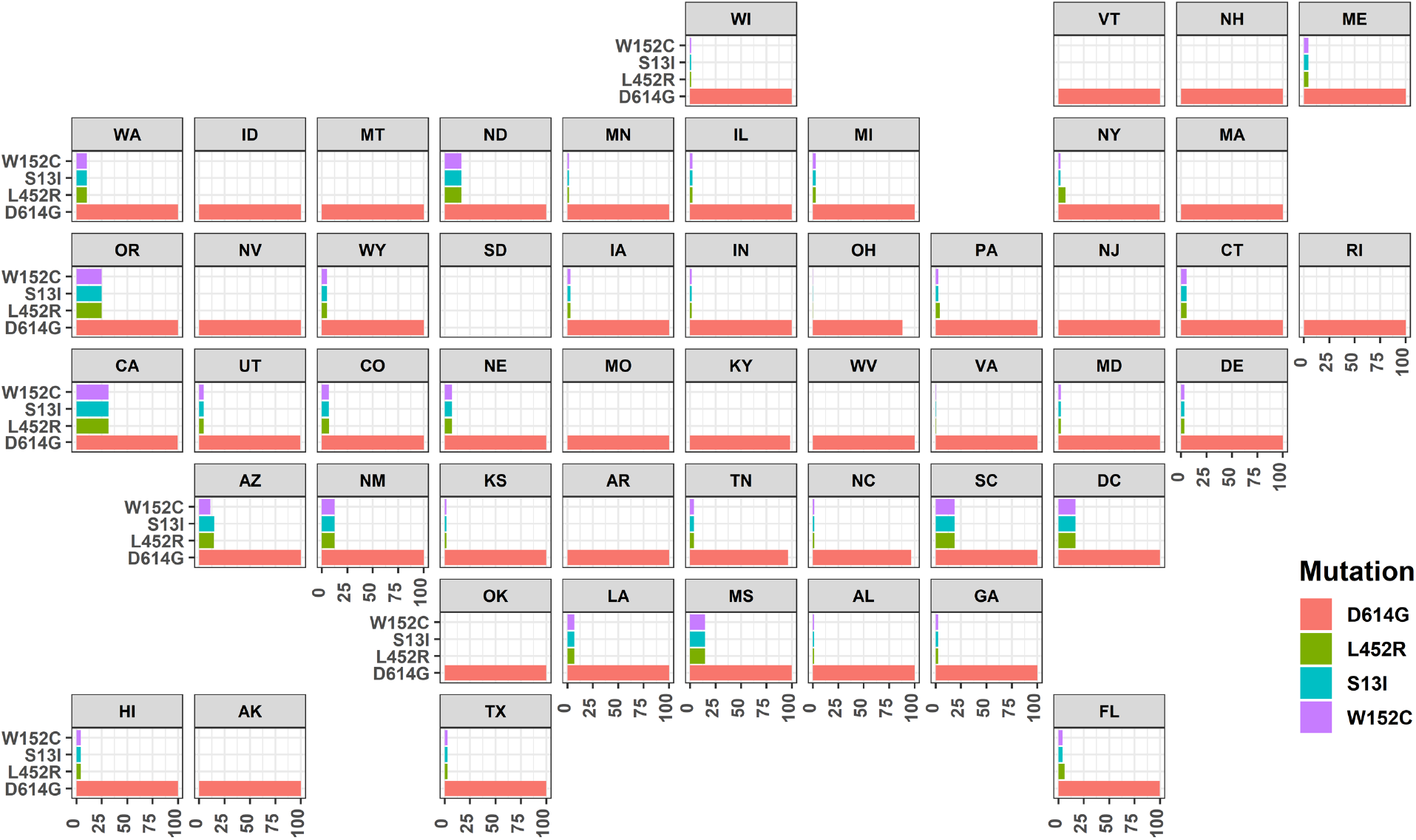
Prevalence of CAL.20C in different states of the United States. Four signature mutations D614G, S13I, W152C and L452R were used to generate this map. This figure was generated by an R programming code that utilizes ‘geofacet’ and ‘ggplot2’ libraries. There were no sequences available from South Dakota. Hence, the mutations could not be plotted.

It is evident that California had the maximum prevalence of CAL.20C (32%) followed by Oregon (25%) > South Carolina (19%) > North Dakota = District of Columbia (17%) > Arizona = Mississippi (15%) > New Mexico (12%) > Washington (10%). The close prevalence of CAL.20C in California and Oregon is not surprising as this variant can easily migrate between these states. However, a 19% prevalence in South Carolina suggests that the virus has migrated directly or through intermediate source from CA to SC during the early emergence of CAL.20C. To our surprise, we did not find CAL.20C mutations from the sequences submitted by Nevada (except D614G). Additionally, a search of GSAID repository showed that the four S protein mutations in that belong to CAL.20C were present in California as early as early July, 2020 (GSAID # hCoV-19/USA/CA-LACPHL-AE00055/2020|EPI_ISL_765994|2020-07-07) [36]. Therefore, the evolution of CAL.20C and its temporal distribution among different states in the U. S. appears rather complex warranting in-depth study.

### 2.8. Structural implications of variant-specific mutations

We used the crystal structure of ACE2/S-RBD (PDB file 6M0J) [9] to understand if the mutations in the S-RBD are topologically positioned such that they enhance the binding of S protein to ACE2. While the effect of some mutations can be directly extrapolated to increased binding, the reported structures of either S protein or the S-RBD/ACE2 complex do not provide direct evidence for the enhanced binding affinity of S protein to the ACE2 receptor. Thus, the signature mutation N501Y, which is present in three of the four variants, forms a hydrophobic interaction between Y501 with Y41 of ACE2 and Y449 of RBD (**Fig. 5a**) that is absent in the case of N501. However, a rationale for greater infectivity of variants with L452R is not obvious from the structural analysis of the reported ACE2/S-RBD complexes. As shown in **Fig. 5b**, L452 does not interact with ACE2, and the mutation R452 in the S-RBD remains too distant to interact with ACE2.

**Figure 5.**
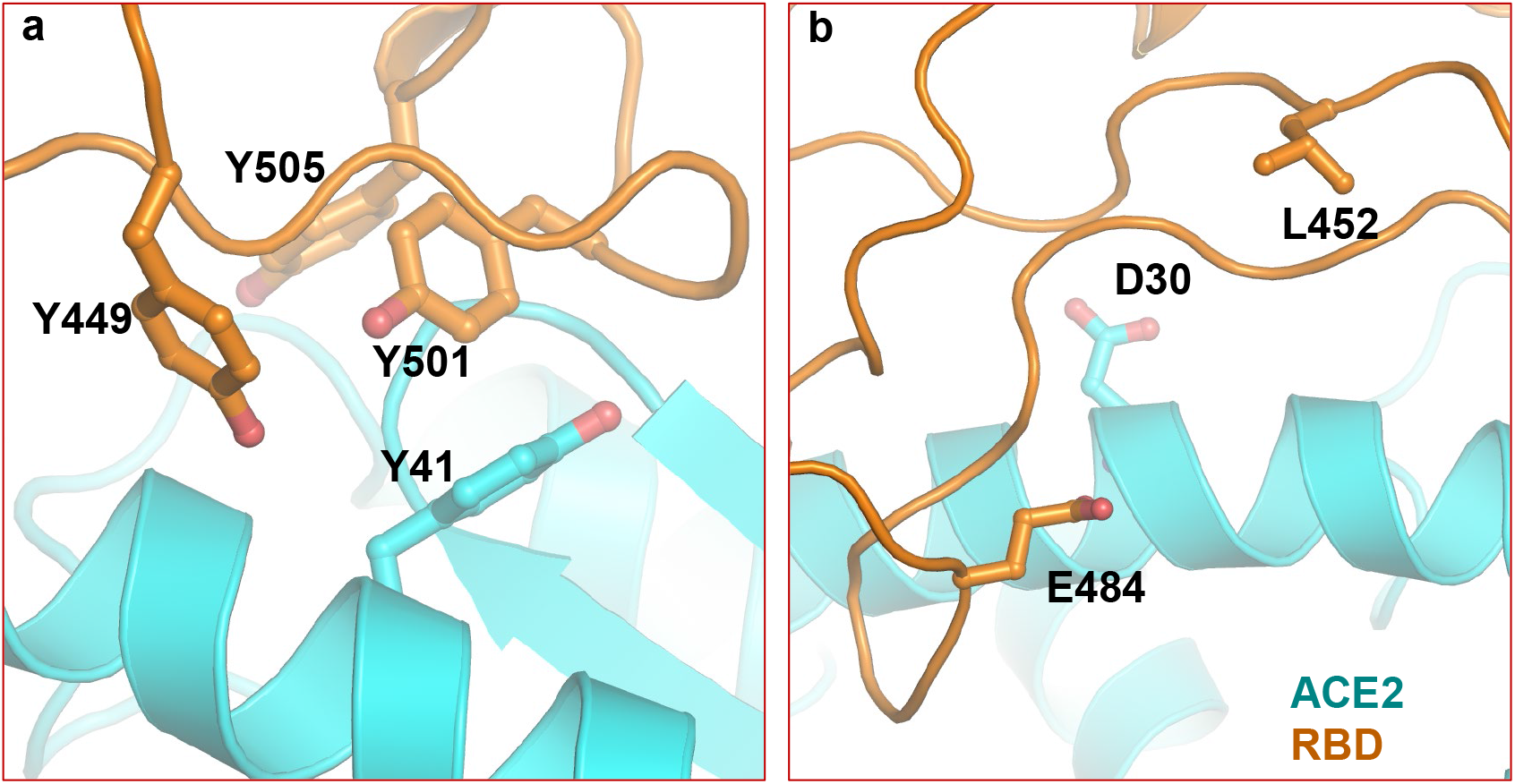
Structural implications of mutations in different variants. This figure depicts two examples of mutations that may enhance the binding of S protein and ACE2 (panel a) or have no effect (panel b).

To gain additional insight, we calculated the change in free energy (ΔΔG) between the wild-type and mutant protein as described by Pandurangan *et al*. [37]. The calculated ΔΔG values for a majority of the signature mutations is shown in **Fig. 6**. A positive value of ΔΔG indicates an increase in protein stability, whereas a negative ΔΔG indicates a decrease in protein stability. Evidently, D614G (the globally dominant virus variant) has the highest ΔΔG, suggesting that 614G S protein is more stable than 614D S protein. K417N, part of the B.1.351 variant, has the lowest ΔΔG, which indicates a decrease in the protein stability. While the computation of ΔΔG does not provide a definitive answer regarding the overall stability of S protein, it does indicate an increased stability of S protein with specific mutations.

**Figure 6.**
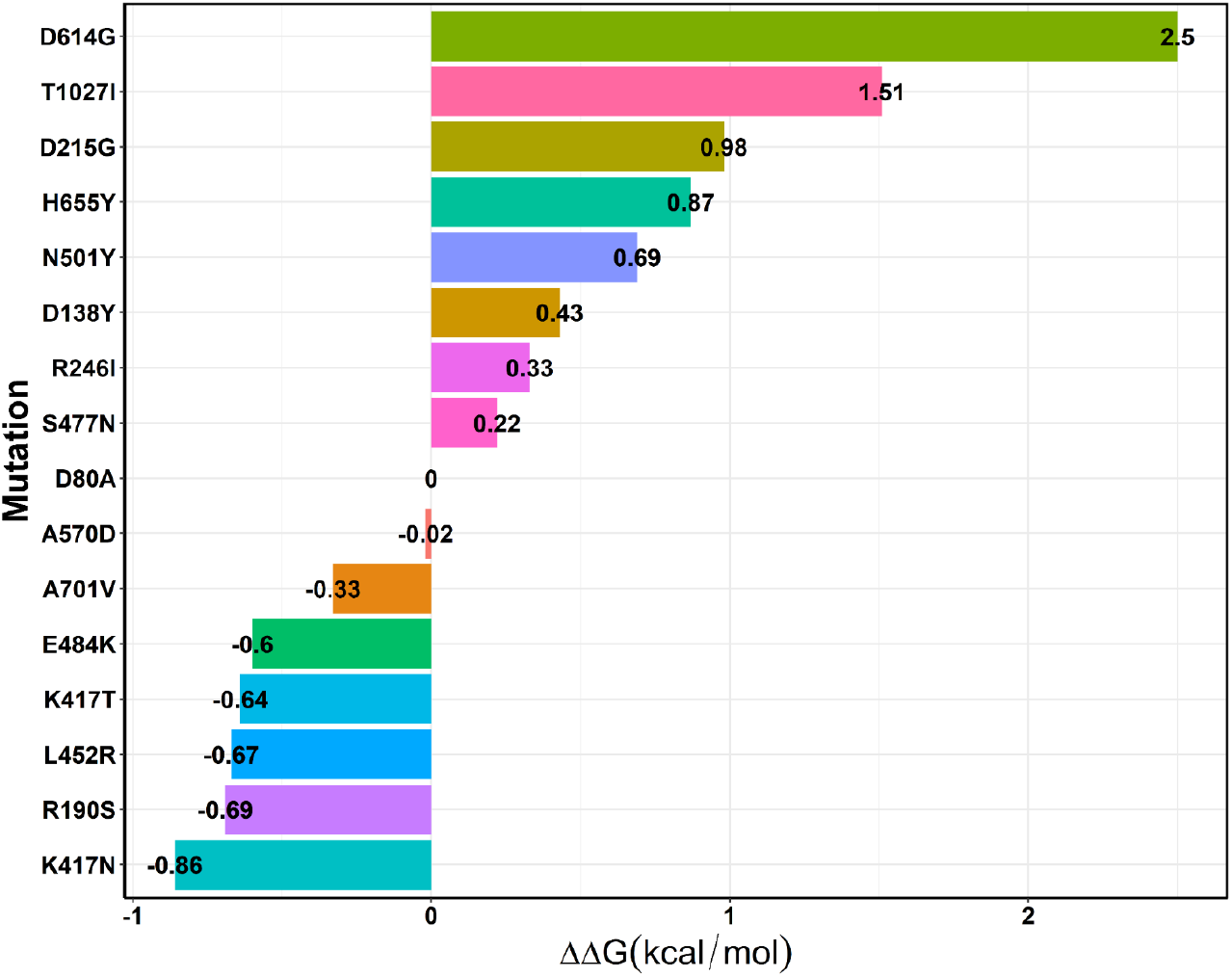
Computed free energy change (ΔΔG) upon mutation. PDB file 6XM4 was used for ΔΔG computation except for N501Y, S477N and E484K, for which PDB file 6M0J [9] was used.

## 3. Summary and Conclusion

Here we present the analyses of factors that may play a role in the higher infectivity of SARS-CoV-2 compared to SARS-CoV. Additionally, we present in-depth analyses of mutations and their prevalence in new SARS-CoV-2 variants that are presumably more infectious than the original (Wuhan-Hu-1) virus (GenBank accession # NC_045512).

The extent of genetic variation between SARS-CoV and SARS-CoV-2 varies depending upon where they circulate in the world. Thus, the nsp coding regions and S protein-coding regions of the two viruses share ∼86% and 76% homology, respectively [38]. The least homologous region in the S protein is the RBM (**Fig. 1a**). However, many amino acid residues within the RBM that participate in ACE2 binding are conserved, and there are only subtle differences in the interaction patterns of SARS-CoV and SARS-CoV-2 S-RBDs. Hence, it is not likely that binding of SARS-CoV and SARS-CoV-2 to ACE2 is the sole determinant influencing infectivity, suggesting that RGD-binding integrin recognition by the RGD motif in the S protein of SARS-CoV-2 may act as an alternate receptor or a co-receptor with ACE2 to enhance viral entry into the host cell. It has been suggested that integrins can bind ACE2 in RGD-dependent and RGD-independent manners, suggesting that ACE2/integrin complexes may provide increased binding opportunities for SARS-CoV-2 with the host cell. Furthermore, integrins present on the cell surface in at least three different conformations: 1) bent, 2) extended conformation with a closed headpiece, and 3) an extended conformation with an open headpiece. These different forms are known to influence the binding of the integrin to its cognate ligands. Because integrins are involved in the recruitment and retention of cells within the inflamed tissues [20,28,39], such as during viral infection, blocking or targeting integrins and the SARS-CoV-2 RBD domain may reduce SARS-COV-2 mediated inflammation or severity of disease, which needs to be further explored in future studies. An unanswered question is compared to the RGD motif in SARS-CoV-2 why do the same ACE2/integrin complexes not confer high SARS-CoV infectivity based upon the presence of the KGD motif? One possibility is that RGD may be recognized by a more significant number of RGD-binding integrins than the KGD motif [40].

Based on the analyses of available sequences of regions of new SARS-CoV-2 variants, it is obvious that these variants have evolved independently. Our correlation analyses of variant-specific mutations also show that the mutations in a given variant may not have evolved simultaneously. As with all RNA viruses, a (resultant) evolved virus goes through quasispecies [41] in which the viral populations consist of mutant spectra (or mutant clouds) rather than genomes with the same nucleotide sequence. It is possible that SARS-CoV-2 genomic sequences rapidly expand in sequence space and lose biological information that enables the elimination of fitness compromised genomes. Subsequent mutations or mutant spectra in the quasispecies of SARS-CoV-2 are most likely generated to circumvent reduced viral fitness for adaptability. This is because the quasispecies mutations constitute dynamic (continuously changing) repositories of genotypic and phenotypic viral variants [41].

From currently available structural, genetic and biochemical data, the higher infectivity of SARS-CoV-2 vs. SARS-CoV is not fully understood, at least in the context of SARS-CoV-2 entry. It would be ideal if a single, relatively discrete host/pathogen interface was primarily responsible for conferring infectivity from a therapeutic perspective. However, our structural studies suggest that it is highly likely that this process is more complex and nuanced. Additional structural information, clinical studies, and a more thorough understanding of the viral life cycle will likely reveal novel biological information and, in turn, illuminate potential therapeutic targets to disrupt SARS-CoV-2 entry.

## 4. Material and Methods

The prevalence of each mutation, in Brazil, California, South Africa and the United Kingdom were determined from the GISAID repository [36]. These sequences were aligned using the MAFFT [42], MEGA [43] or JalView [44] sequence alignment programs. The sequences were then analyzed through in-house Python or R scripts. Relative abundance (RA) of mutations in S protein from different geographic regions was determined by an in-house python script using the scikit-learn (Python) library [45] and plotted with R (codes available upon request). Final values were multiplied by 100 to express as a percent. For the correlation analyses, an R package ‘ggcorrplot’ was used in an in-house R script. The prevalence of CAL.20C within the United States was computed by an in-house Python script. All in-house scripts are available upon request. The structural analyses were conducted using the Schrodinger Suite (Schrodinger LLC, NY) and PyMol [46]. The free energy change upon mutation in S protein (ΔΔG) was computed through SDM server as detailed by Pandurangan *et al*. [37].

## 5. Author contributions

KS, ANS and SRK conceptualized the study; KS wrote the first draft. ANS, SRK and KS conducted genetic analyses; LTW, GAW and SNB contributed to defining the integrin relationship to SARS-CoV-2 infectivity and editing the manuscript; TPQ, AS and CLL edited the manuscript and contributed to understanding the pathogenicity of CoVs. All authors approved the final manuscript.

## 6. Acknowledgments

KS acknowledges support from the Office of Research, University of Missouri (Bond Life Sciences Center, Early Concept Grant). AS acknowledges funding from the Swedish Research Council (2016-01675). LTW, GAW and KS acknowledge support from the National Institute of Dental and Craniofacial Research Grant DE007389 and the American Lung Association. KS acknowledges National Institute of General Medicine Grant GM118012. KS and TPQ acknowledge the computation facilities of the Molecular Interaction Core at the University of Missouri, Columbia, MO 65212.

---

**Supplementary Fig. S1:**
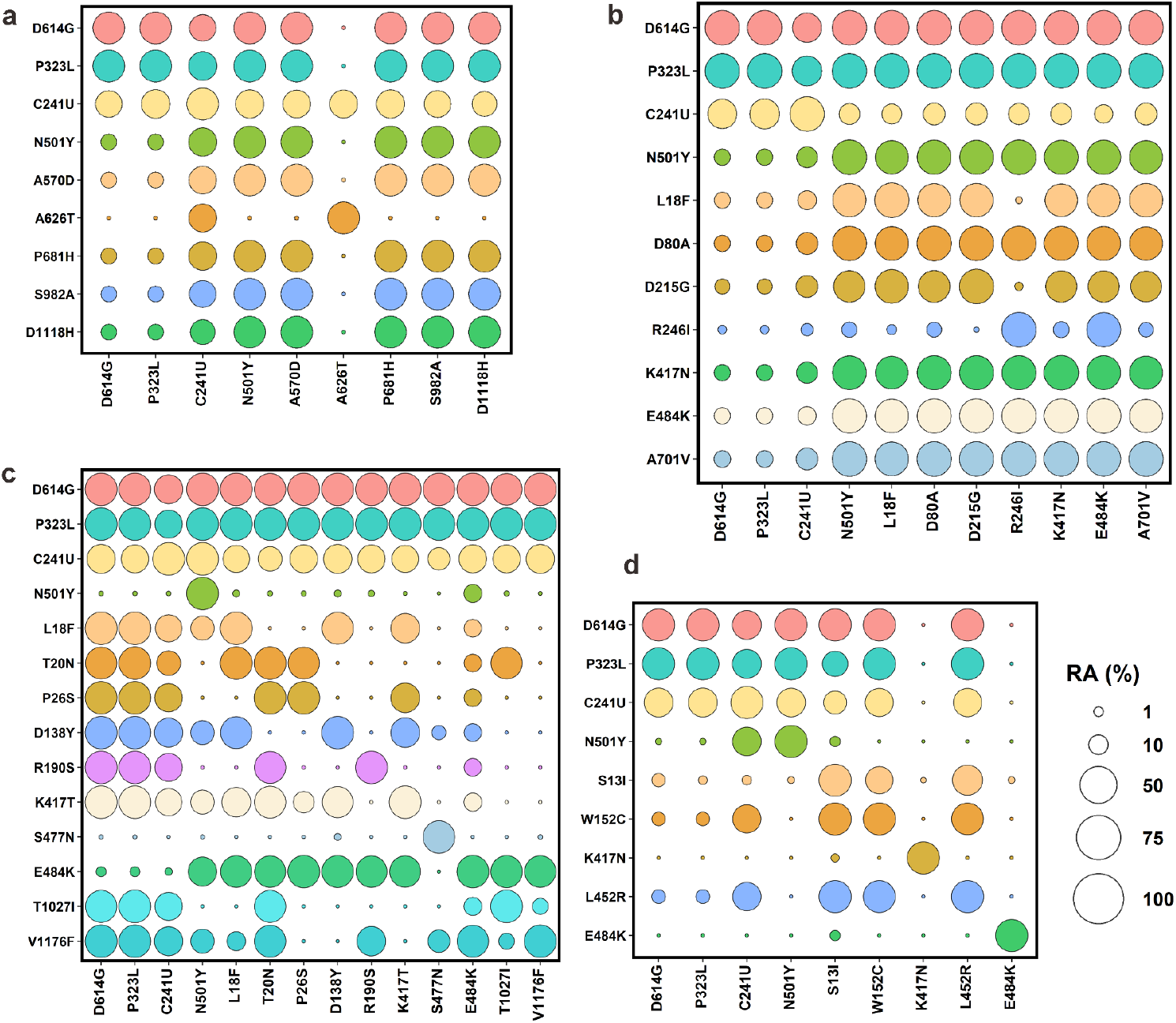
Relative abundance (RA) of mutations in S protein. Panels a, b, c and d show the RA of mutations in variants B.1.1.7 (UK), B1.351 (South Africa), P.1 (Brazil) and CAL.20C (California), respectively. The normalized RA was calculated using an in-house python script that utilized the scikit-learn (Python) library. The figure was made with R (codes available upon request).

**Supplementary Fig. S2:**
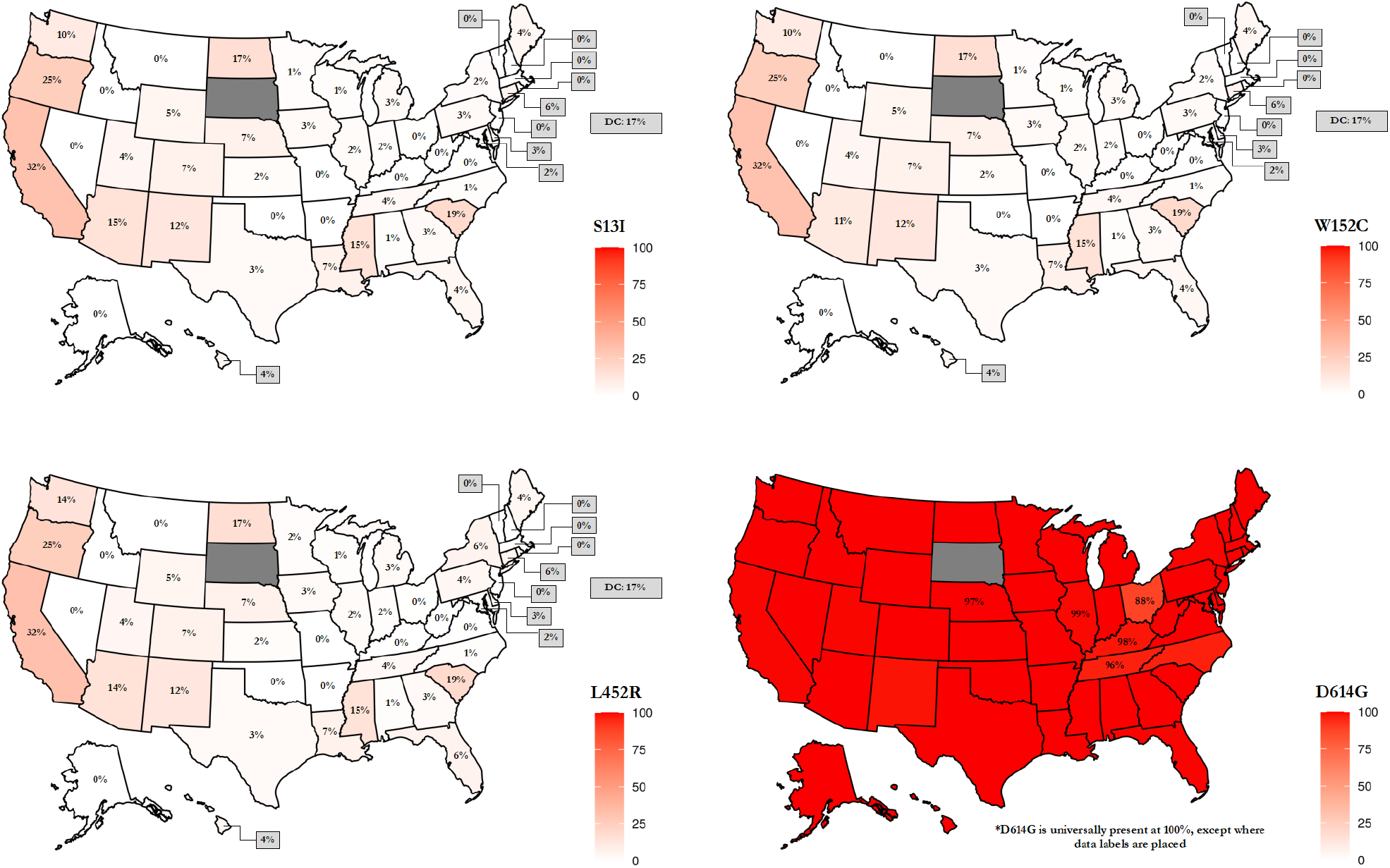
Prevalence of CAL.20C in different states of the U.S.

## SUPPLEMENTARY TEXT

### Known mutations (in addition to those in S protein) in different variants

#### B.1.1.7 Lineage (Britain)

1. N protein
  a. S235F mutation
  b. D3L
  c. Y73C
  d. R521
2. ORF8 protein
  a. **Q27 stop** (Changes 27^th^ amino acid from Q to STOP codon and leaves a 26 amino acid long stump; 2 other mutations appear after the stop but since the protein is cut short, they may have no function)
3. ORF1a protein
  a. SGF 3675-3677 deletion
  b. I2230T
  c. A1708D
  d. T1001I
4. Six additional silent mutations

#### B.1.351 Lineage (South Africa)

1. N protein
  a. T205I mutation
2. M/E
  a. P71L
3. ORF1a protein
  a. SGF 3675-3677 deletion
  b. K1655N

#### P.1 Lineage (Japan > Brazil)

1. N protein
  a. P80R mutation
  b. 28269-73 insertion
  c. E92K
2. ORF1b protein
  a. E5665D
3. ORF1a protein
  a. SGF 3675-3677 deletion
  b. K1795Q
  c. S1188L

#### CAL.20C (California)

1. ORF1a protein
  a. I4205V
2. ORF1b protein
  a. D1183Y

## References

1. Kim, D., Lee, J.Y., Yang, J.S., Kim, J.W., Kim, V.N., Chang, H. The architecture of SARS-CoV-2 transcriptome. Cell 2020.

2. Snijder, E.J., Bredenbeek, P.J., Dobbe, J.C., Thiel, V., Ziebuhr, J., Poon, L.L., Guan, Y., Rozanov, M., Spaan, W.J., Gorbalenya, A.E. Unique and conserved features of genome and proteome of SARS-coronavirus, an early split-off from the coronavirus group 2 lineage. J Mol Biol 2003, 331, 991–1004.

3. Masters, P.S. The molecular biology of coronaviruses. Adv Virus Res 2006, 66, 193–292.

4. Ziebuhr, J. Molecular biology of severe acute respiratory syndrome coronavirus. Curr Opin Microbiol 2004, 7, 412–419.

5. Yan, L., Ge, J., Zheng, L., Zhang, Y., Gao, Y., Wang, T., Huang, Y., Yang, Y., Gao, S., Li, M., Liu, Z., Wang, H., Li, Y., Chen, Y., Guddat, L.W., Wang, Q., Rao, Z., Lou, Z. Cryo-EM structure of an extended SARS-CoV-2 replication and transcription complex reveals an intermediate state in cap synthesis. Cell 2021, 184, 184–193 e110.

6. Dai, L., Gao, G.F. Viral targets for vaccines against COVID-19. Nat Rev Immunol 2021, 21, 73–82.

7. Walls, A.C., Park, Y.J., Tortorici, M.A., Wall, A., McGuire, A.T., Veesler, D. Structure, function, and antigenicity of the SARS-CoV-2 spike glycoprotein. Cell 2020, 181, 281–292 e286.

8. Wrapp, D., Wang, N., Corbett, K.S., Goldsmith, J.A., Hsieh, C.L., Abiona, O., Graham, B.S., McLellan, J.S. Cryo-EM structure of the 2019-nCoV spike in the prefusion conformation. Science 2020, 367, 1260–1263.

9. Lan, J., Ge, J., Yu, J., Shan, S., Zhou, H., Fan, S., Zhang, Q., Shi, X., Wang, Q., Zhang, L., Wang, X. Structure of the SARS-CoV-2 spike receptor-binding domain bound to the ACE2 receptor. Nature 2020, 581, 215–220.

10. Li, F., Li, W., Farzan, M., Harrison, S.C. Structure of SARS coronavirus spike receptor-binding domain complexed with receptor. Science 2005, 309, 1864–1868.

11. Wu, K., Li, W., Peng, G., Li, F. Crystal structure of NL63 respiratory coronavirus receptor-binding domain complexed with its human receptor. Proc Natl Acad Sci U S A 2009, 106, 19970–19974.

12. Wang, Q., Zhang, Y., Wu, L., Niu, S., Song, C., Zhang, Z., Lu, G., Qiao, C., Hu, Y., Yuen, K.Y., Wang, Q., Zhou, H., Yan, J., Qi, J. Structural and functional basis of SARS-CoV-2 entry by using human ACE2. Cell 2020, 181, 894–904 e899.

13. Shang, J., Ye, G., Shi, K., Wan, Y., Luo, C., Aihara, H., Geng, Q., Auerbach, A., Li, F. Structural basis of receptor recognition by SARS-CoV-2. Nature 2020, 581, 221–224.

14. Yan, R., Zhang, Y., Li, Y., Xia, L., Guo, Y., Zhou, Q. Structural basis for the recognition of SARS-CoV-2 by full-length human ACE2. Science 2020, 367, 1444–1448.

15. Benton, D.J., Wrobel, A.G., Xu, P., Roustan, C., Martin, S.R., Rosenthal, P.B., Skehel, J.J., Gamblin, S.J. Receptor binding and priming of the spike protein of SARS-CoV-2 for membrane fusion. Nature 2020, 588, 327–330.

16. Walls, A.C., Park, Y.-J., Tortorici, M.A., Wall, A., McGuire, A.T., Veesler, D. Structure, function, and antigenicity of the SARS-CoV-2 spike glycoprotein. Cell 2020, 181, 281–292.e286.

17. Sigrist, C.J., Bridge, A., Le Mercier, P. A potential role for integrins in host cell entry by SARS-CoV-2. Antiviral Res 2020, 177, 104759.

18. Scarborough, R.M., Rose, J.W., Hsu, M.A., Phillips, D.R., Fried, V.A., Campbell, A.M., Nannizzi, L., Charo, I.F. Barbourin. A gpiib-iiia-specific integrin antagonist from the venom of sistrurus m. Barbouri. J Biol Chem 1991, 266, 9359–9362.

19. Barczyk, M., Carracedo, S., Gullberg, D. Integrins. Cell Tissue Res 2010, 339, 269–280.

20. Humphries, J.D., Byron, A., Humphries, M.J. Integrin ligands at a glance. J Cell Sci 2006, 119, 3901–3903.

21. Bagchi, S., Liao, Z., Gonzalez, F.A., Chorna, N.E., Seye, C.I., Weisman, G.A., Erb, L. The p2y2 nucleotide receptor interacts with alphav integrins to activate go and induce cell migration. J Biol Chem 2005, 280, 39050–39057.

22. Erb, L., Liu, J., Ockerhausen, J., Kong, Q., Garrad, R.C., Griffin, K., Neal, C., Krugh, B., Santiago-Perez, L.I., Gonzalez, F.A., Gresham, H.D., Turner, J.T., Weisman, G.A. An rgd sequence in the p2y(2) receptor interacts with alpha(v)beta(3) integrins and is required for g(o)-mediated signal transduction. J Cell Biol 2001, 153, 491–501.

23. Liao, Z., Seye, C.I., Weisman, G.A., Erb, L. The p2y2 nucleotide receptor requires interaction with alpha v integrins to access and activate g12. J Cell Sci 2007, 120, 1654–1662.

24. Stewart, P.L., Nemerow, G.R. Cell integrins: Commonly used receptors for diverse viral pathogens. Trends Microbiol 2007, 15, 500–507.

25. Wei, Y., Zhang, Y., Cai, H., Mirza, A.M., Iorio, R.M., Peeples, M.E., Niewiesk, S., Li, J. Roles of the putative integrin-binding motif of the human metapneumovirus fusion (f) protein in cell-cell fusion, viral infectivity, and pathogenesis. J Virol 2014, 88, 4338–4352.

26. Feire, A.L., Koss, H., Compton, T. Cellular integrins function as entry receptors for human cytomegalovirus via a highly conserved disintegrin-like domain. Proc Natl Acad Sci U S A 2004, 101, 15470–15475.

27. Wickham, T.J., Filardo, E.J., Cheresh, D.A., Nemerow, G.R. Integrin alpha v beta 5 selectively promotes adenovirus mediated cell membrane permeabilization. J Cell Biol 1994, 127, 257–264.

28. Ansari, A.A., Byrareddy, S.N. The role of integrin expressing cells in modulating disease susceptibility and progression (january 2016). Int Trends Immun 2016, 4, 11–27.

29. Clarke, N.E., Fisher, M.J., Porter, K.E., Lambert, D.W., Turner, A.J. Angiotensin converting enzyme (ace) and ACE2 bind integrins and ACE2 regulates integrin signalling. PLoS One 2012, 7, e34747.

30. Li, W., Zhang, C., Sui, J., Kuhn, J.H., Moore, M.J., Luo, S., Wong, S.K., Huang, I.C., Xu, K., Vasilieva, N., Murakami, A., He, Y., Marasco, W.A., Guan, Y., Choe, H., Farzan, M. Receptor and viral determinants of SARS-coronavirus adaptation to human ACE2. EMBO J 2005, 24, 1634–1643.

31. Danilczyk, U., Sarao, R., Remy, C., Benabbas, C., Stange, G., Richter, A., Arya, S., Pospisilik, J.A., Singer, D., Camargo, S.M., Makrides, V., Ramadan, T., Verrey, F., Wagner, C.A., Penninger, J.M. Essential role for collectrin in renal amino acid transport. Nature 2006, 444, 1088–1091.

32. Malakauskas, S.M., Quan, H., Fields, T.A., McCall, S.J., Yu, M.J., Kourany, W.M., Frey, C.W., Le, T.H. Aminoaciduria and altered renal expression of luminal amino acid transporters in mice lacking novel gene collectrin. Am J Physiol] Renal Physiol 2007, 292, F533–544.

33. Kowalczuk, S., Broer, A., Tietze, N., Vanslambrouck, J.M., Rasko, J.E., Broer, S. A protein complex in the brush-border membrane explains a hartnup disorder allele. FASEB J 2008, 22, 2880–2887.

34. Fairweather, S.J., Broer, A., O’ Mara, M.L., Broer, S. Intestinal peptidases form functional complexes with the neutral amino acid transporter b(0)at1. Biochem J 2012, 446, 135–148.

35. Kannan, S.R., Spratt, A.N., Quinn, T.P., Heng, X., Lorson, C.L., Sonnerborg, A., Byrareddy, S.N., Singh, K. Infectivity of SARS-CoV-2: There is something more than D614G? J Neuroimmune Pharmacol 2020, 15, 574–577.

36. Elbe, S., Buckland-Merrett, G. Data, disease and diplomacy: Gisaid’ s innovative contribution to global health. Glob Chall 2017, 1, 33–46.

37. Pandurangan, A.P., Ochoa-Montano, B., Ascher, D.B., Blundell, T.L. SDM: A server for predicting effects of mutations on protein stability. Nucleic Acids Res 2017, 45, W229–W235.

38. Grifoni, A., Sidney, J., Zhang, Y., Scheuermann, R.H., Peters, B., Sette, A. A sequence homology and bioinformatic approach can predict candidate targets for immune responses to SARS-CoV-2. Cell Host Microbe 2020, 27, 671–680 e672.

39. Humphries, M.J. Integrin structure. Biochem Soc Trans 2000, 28, 311–339.

40. Luan, J., Lu, Y., Gao, S., Zhang, L. A potential inhibitory role for integrin in the receptor targeting of SARS-CoV-2. J Infect 2020.

41. Domingo, E., Sheldon, J., Perales, C. Viral quasispecies evolution. Microbiol Mol Biol Rev 2012, 76, 159–216.

42. Katoh, K., Standley, D.M. MAFFT multiple sequence alignment software version 7: Improvements in performance and usability. Mol Biol Evol 2013, 30, 772–780.

43. Kumar, S., Stecher, G., Li, M., Knyaz, C., Tamura, K. MEGA x: Molecular evolutionary genetics analysis across computing platforms. Mol Biol Evol 2018, 35, 1547–1549.

44. Troshin, P.V., Procter, J.B., Barton, G.J. Java bioinformatics analysis web services for multiple sequence alignment--jabaws:MSA. Bioinformatics 2011, 27, 2001–2002.

45. Pedregosa, F., Varoquaux, G., Gramfort, A., Michel, V., Thirion, B., Grisel, O., Blondel, M., Prettenhofer, P., Weiss, R., Dubourg, V., Vanderplas, J., Passos, A., Cournapeau, D., Brucher, M., Perrot, M., Duchesnay, E. Scikit-learn: Machine learning in python. Journal of Machine Learning Research 2011, 12, 2825–2830.

46. DeLano, W.L. An open-source molecular graphics tool. CCP4 newsletter on protein crystallography. 2002, 40, 82–92.

